# Longitudinal examination of perfusion and angiogenesis markers in primary colorectal tumors shows distinct signatures for metronomic and maximum-tolerated dose strategies

**DOI:** 10.1101/2022.02.07.479423

**Authors:** Ariel I. Mundo, Abdussaboor Muhammad, Kerlin Balza, Christopher E. Nelson, Timothy J. Muldoon

**Affiliations:** Department of Biomedical Engineering, University of Arkansas, Fayetteville, AR, USA

**Keywords:** longitudinal data, colon cancer, azoxymethane, hypoxia, vegf, generalized additive models

## Abstract

Metronomic chemotherapy (MET) has been developed to address the shortcomings of maximum-tolerated chemotherapy (MTD) in regard to toxicity and development of resistance mechanisms in the tumor. In colorectal cancer (CRC), MET is a promising novel strategy to treat locally advanced malignancies when used as neoadjuvant chemotherapy (NAC). However, so far there are no preclinical studies to assess the impact of MET NAC in CRC to assess the benefits and challenges of this approach. Here, we used a primary model of CRC (via azoxymethane) to analyze longitudinal changes in angiogenesis in primary tumors under MET and MTD NAC using a combination of diffuse reflectance spectroscopy and mRNA expression (via qPCR). Our results show that MET and MTD NAC lead to increased mean tissue oxygen saturation (8% and 5%, respectively) and oxyhemoglobin (15% and 10%) between weeks 2 and 5 of NAC, and that such increases are caused by distinct molecular signatures in the angiogenic program. Specifically, we find that in the MET group there is a sustained increase in *Hif-1a, Aldoa*, and *Pgk1* expression, suggesting upregulated glycolysis, whereas MTD NAC causes a significant reduction in the expression of the aforementioned genes and of *Vegf*, leading to vascular remodeling in MTD-treated tumors. Taken together, this study demonstrates the ability of combined optical and molecular methodologies to provide a holistic picture of tumor response to therapy in CRC in a minimally invasive manner.

## 2 Introduction

Colorectal cancer (CRC) is the third most commonly diagnosed cancer in the US^1^. Despite an overall decline in over the last two decades, it is known that individuals younger than 50 years^2^ have shown an increasing trend in incidence, which is likely to hold in the near future. The majority of patients (about 75%, regardless of age) present at diagnosis what is known as “locally advanced disease”. Locally advanced disease is a term used to group malignancies that range from those tumors that have only invaded the colonic mucosa to those that have spread to adjacent lymph nodes(stages IIA-IIIC)^3^.

In biomedical research, there is an active area of interest in novel treatment strategies that can be used to treat locally advanced disease in CRC. In this regard, an increasing body of evidence has shown that neoadjuvant chemotherapy (defined as chemotherapy administered to treat tumors before surgical resection) is able to reduce tumor size, facilitate resection, and improve completion rates in CRC^4–7^. The standard of care in neoadjuvant chemotherapy (NAC) consists in the use of maximum-tolerated doses (MTDs) of a cytotoxic drug. For CRC, the fluoropyrimidine 5-fluorouracil (5-FU) is the most commonly used cytotoxic agent used either alone or in combination with other drugs that aid its therapeutic effects^8^.

Although the goal of MTD NAC is to reduce tumor burden as much as possible, its associated toxicity makes necessary the delivery of therapy in a cyclic manner. For example, the FOLFOX regimen (a combination of leucovorin, 5-FU and oxaliplatin) is delivered in cycles that last two weeks; only the first 48 hours are of active infusion, followed by 12 days of rest^9,10^. Because a complete MTD NAC treatment may last between 3-4 cycles (6-8 weeks)^11^, the rest periods between each cycle present an opportunity for the tumor to recover and develop resistance. Indeed, it has been shown that MTD NAC is associated with re-growth of tumor cells and the development of drug resistance^12,13^. Altogether, the shortcomings of MTD NAC have demonstrated the need of new therapy strategies that can effectively reduce tumor burden while minimizing the associated undesirable side-effects caused by high-dose chemotherapy.

During the last decade, the biological effect of chemotherapeutic drugs at doses below the MTD threshold started to be explored. The term “metronomic chemotherapy” was coined to describe chemotherapy that involves the use of lower but active doses of chemotherapeutic drugs without prolonged rest periods in between^14^. Early studies on metronomic chemotherapy (MET), showed that the targets of this type of treatment were the endothelial cells of the tumor, which could not recover due to the continuous delivery of the drugs^14,15^, which translated in an anti-angiogenic effect^16^.

Various studies have explored the use of MET in recurring and late-stage malignancies such as colon, glioma, breast, and ovarian cancer where it has been shown to improve survival^17,18^, reduce toxicity^19^, and cause high pathological complete response (no residual invasive disease on pathological evaluation)^20^. However, the potential use of MET as an effective alternative to MTD NAC in CRC has not been examined so far.

More specifically, although it is assumed that the main effect of MET is its ability to disrupt the angiogenic program of tumors by impairing blood vessel development, it is not known how this effect varies between different types of malignancies. In particular, it is possible that the anti-angiogenic effect of MET is heavily dependent of the anatomic location of the tumor, as it is known from preclinical studies that the tumor milieu plays a critical role in the response to treatment^21,22^. For example, in the case of malignancies that arise in a lipid-rich environment (such as the colon), it has been shown that a hypoxia-driven shift in lipid metabolism can be coupled with tumor growth without the presence of a dense vascular network^23^. Therefore, there is a need for an improved understanding of the metabolic implications of MET NAC in CRC in regard to angiogenesis, and how these changes contrast with those elicited by MTD NAC.

In this study, we hypothesized that the differences in dosing and frequency of administration between MTD and MET NAC would cause distinct changes in the angiogenic program of primary colorectal tumors that could be analyzed to better understand the opportunities and pitfalls of MET NAC when used on CRC. To this end, we used a dual approach to examine ‘high level’ and ‘low level’ changes in angiogenesis. Firstly, we used endoscopic diffuse reflectance spectroscopy (DRS), an optical technique that enables quantification of physiological metrics of perfusion in tissue such as oxygen saturation (StO_2_), oxyhemoglobin (HbO_2_), and deoxyhemoglobin (HbO), to study the ‘high level’ effect of therapy in tumor perfusion. Because DRS is non-destructive, and relatively low-cost, it is useful to obtain an overall assessment of tumor perfusion and response to therapy^24,25^.

Secondly, we also examined the genes (via mRNA expression) involved in the classical hypoxia-inducible factor 1 (HIF-1A) and vascular endothelial growth factor (VEGF) angiogenic axis to examine molecular (‘low level’) changes that could provide the rationale for the observed changes in perfusion obtained via DRS. In additionh, gene expression data were used to gain insight in the metabolic alterations that therapy progression causes in the tumor for both MET and MTD NAC. Our study addresses the need to connect ‘high level’ reporters of perfusion of DRS with molecular changes via qPCR to better understand the challenges and opportunities of MTD and MET NAC strategies in CRC.

## 3 Materials and Methods

### 3.1 Animal model

All procedures were approved by the University of Arkansas Institutional Animal Care and Use Committee under protocol 21024. An azoxymethane (AOM) model of CRC was used where nine-week-old female A/J mice (000646,The Jackson Laboratory) received AOM injections (A5486, Millipore Sigma) at a dose of 10 mg/kg once a week for six weeks^26^. After acclimation for one week upon arrival mice received AOM injections (Figure 1,A); mice were weighed daily and received a combination or rodent chow (8640, Teklad) and hydrogel (Nutra-Gel, AIN-93, Bio-Serv); if weight loss was greater than 10% (compared to the weight on the first day of AOM injections), they received 1500 *μ*L of 0.9% sterile saline solution (S5825, Teknova) via subcutaneous injection.

**Figure 1:**
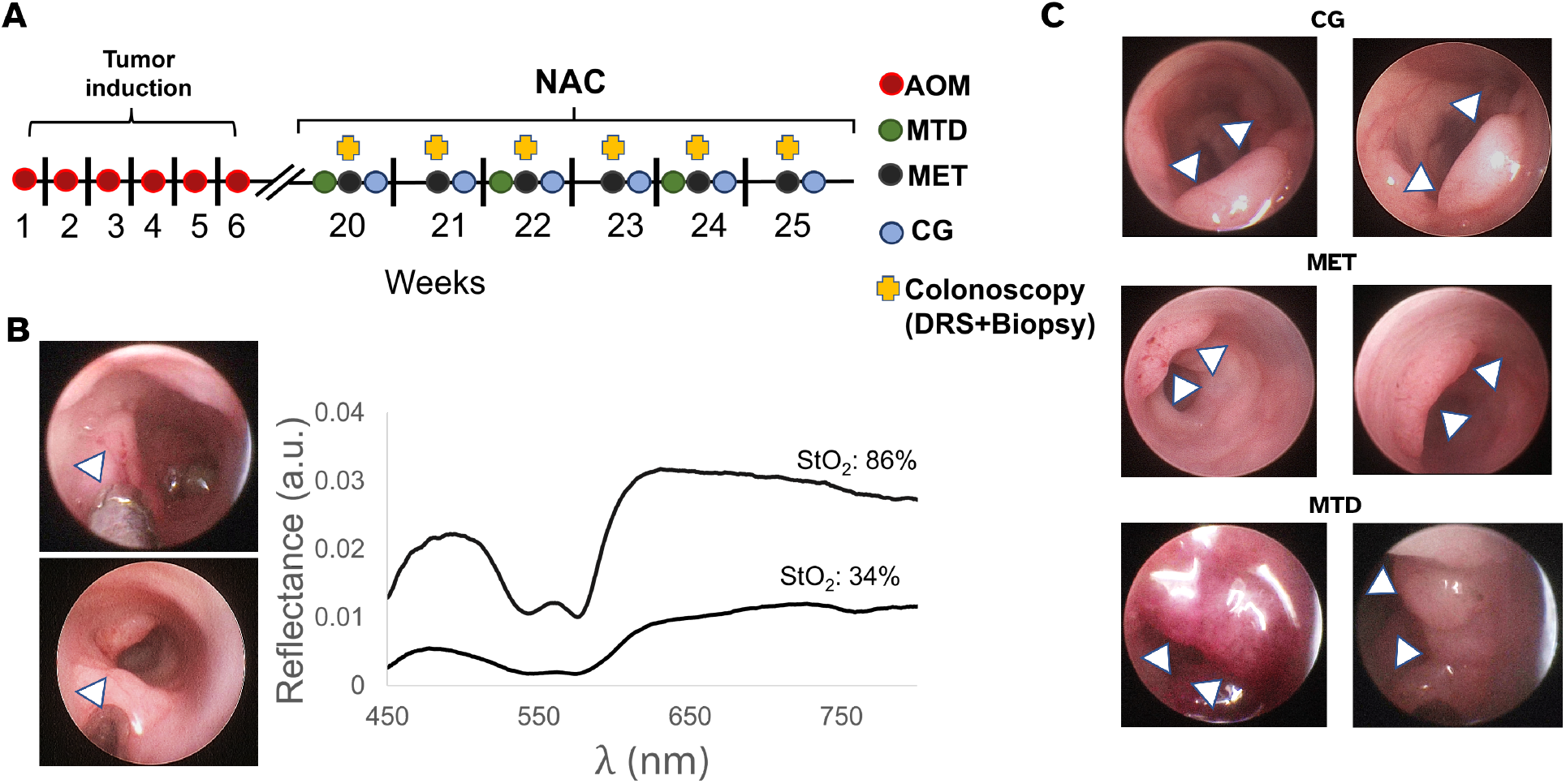
Study design and endoscopically-acquired representative images of primary colorectal tumors. **A**: After AOM induction for six weeks, mice received either MTD NAC (20 mg/kg of 5-FU for five consecutive days every other week), MET NAC (8 mg/kg every other day three times a week), or saline (CG) treatment (200 *μ*L). The next day after the first injection mice started to receive endoscopic examination which included DRS and biopsy acquisition. **B**: Representative images of endoscopic procedures and optically-derived perfusion values. *Top*: Biopsy acquisition using the biopsy forceps (indicated by arrow). *Bottom*: DRS probe (indicated by arrow) placement on a tumor previous to spectra acquisition. DRS probe diameter was ≈ 0.78 mm. *Right*: Representative optically-derived values of StO_2_ obtained after spectra post-processing. Top line shows the reflectance from a tumor showing high StO_2_, note the distinct Q-bands in the reflectance (between 500-575 nm). Bottom line shows reflectance from a hypoxic tumor, note the low StO_2_ value and the flat profile of the reflectance in the 475-600 nm range. **C**: Representative endoscopic images of the changes in tumor size of the course of therapy. *Top*: Tumors from the CG group. *Middle*: Tumors from the MET group. Tumors in the CG and MET groups showed no reduction in size over the course of therapy, their size increasing as therapy progressed (top and middle). Arrows indicate tumor location, note the tumor vasculature which is endoscopically visible. *Bottom*: MTD-trated tumors. These tumors showed a cyclic pattern of tumor growth and shrinkage with every therapy cycle. Note the intense dark red color on the tumor on the left indicating hypoxia, and its change in color and reduction in size in the following colonoscopy (right).

### 3.2 Endoscopic Imaging

At 16 weeks after the last AOM injection the size of tumors was periodically assessed using a veterinary colonoscopy system (ColoVIEW, Karl Storz) as we have previously described^27^. Briefly, mice were anesthetized using 2% isoflurane using a precision vaporizer and placed on a heating pad set at 42°C to maintain body temperature through the procedure. Endoscopic procedures began when mice had no pedal reflex. Fecal matter was removed by flushing the colon with 0.9% sterile saline, and the endoscopic examination consisted in the careful insertion of the endoscopic sheath containing the telescope up to a flexure at approximately 4cm as it has been reported elsewhere^28^. From there, the examination sheath was slowly retracted from the colon while acquiring imaging data (pictures, video) to determine tumor location; tumors were labelled sequentially following the proximal to distal orientation and their size was graded relative to the circumference of the colon^29^. Once tumors were determined to be ~ 1mm in diameter mice were randomly distributed in different chemotherapy groups.

### 3.3 Chemotherapy

Chemotherapy was based on 5-FU (F6627, Millipore Sigma), which was first solubilized in dimethyl sulfoxide (97063-136, VWR) to a concentration of 50 mg/mL. This solution was then diluted in sterile saline to concentrations of 3.0 mg/mL (for the MTD group) and of 1.2 mg/mL (for the MET group), stored at −20°C and thawed before administration. Mice were randomly distributed between maximum-tolerated (MTD, n=27), metronomic (MET, n=20) and control (CG, n=14) groups. All therapy was provided via intraperitoneal injection as follows: MTD, 20 mg/kg of 5-FU for five consecutive days every other week; MET, 8 mg/kg of 5-FU every other day three times a week; CG, 200 *μ*L of sterile saline every other day (Figure 1A). Dosing and schedule for administration were determined using both reported MTD of 5-FU MTD in mice^30^, and the NAC cycling in the clinical setting. For MET, we followed previously reported MET 5-FU dosing in mice^31,32^. The next day of the first treatment injection (either 5-FU or saline), mice started to receive endoscopic examination for the acquisition of DRS and biopsies.

### 3.4 Endoscopic DRS and biopsy acquisition

The next day of the first 5-FU (or saline) injection, mice underwent a colonoscopy as previously described and DRS spectra were acquired as we have reported elsewhere^27^. Briefly, DRS were acquired using a USB spectrometer (Flame-T, Ocean Insight), and a sub-millimeter probe (Myriad Fiber Imaging, Dudley, MA, US). The probe diameter was ≈ 0.78 mm, with ≈ 290*μ*m SDS and was connected to the insufflation port of the endoscopic examination sheath while using a shutter to block the light from the colonoscopy system lamp to avoid contamination on the spectra. The light source for DRS acquisition was a tungsten halogen lamp (HL-2000, Ocean Insight) whose intensity was calibrated using a 5% reflectance standard (SRS-05-010, Labsphere). All spectra were collected using an integration time of 75 ms.

Spectra were post-processed to obtain perfusion values from tissue (StO_2_, HbO_2_, etc.) as we have described in detail elsewhere^27^. Briefly, we used an inverse lookup table (LUT) model^33^ to extract absorption and scattering properties from tumors from spectra which were used to obtain perfusion values. We used a custom algorithm to discard spectra with artifacts from operator movement, mouse breathing or excessive pressure at the tip of the probe^27^. After DRS spectra were acquired, the endoscopic sheath was cleaned with 70% ethanol and RNAse Away (7002, ThermoFisher), and a biopsy from the tumor was obtained using a 3 Fr. biopsy forceps (61071 ZJ, Karl Storz). Biopsies were placed in RNALater (AM7021, ThermoFisher) using a 0.5 sterile microcentrifuge tube (022600001, Eppendorf) and stored at −80°C. Biopsies for the MET and CG group were obtained at weeks 1, 4 and 6, whereas in the MTD group biopsies were acquired weekly. After biopsy acquisition mice were removed from anesthesia once no active bleeding from the biopsy site was determined. Endoscopies were repeated once every 7 days for a total of 6 colonoscopies for every mouse.

### 3.5 Quantitative Real-Time PCR

Biopsies were homogenized over ice using a sonicator (Q55 Sonicator, QSonica), and total mRNA extraction was performed using the RNAeasy Micro Kit (74004, Qiagen) following manufacturer recommendations. Quality of the extracted mRNA was determined using a spectrophotometer (NanoDrop One, ThermoFisher), using samples whose 260/280 absorbance ratio ranged between 1.8-2.0^34,35^. Reverse transcription to cDNA was performed using the SuperScript Synthesis VILO cDNA Kit (ThermoFisher, 11754050), and cDNA concentration was normalized at 2.5 ng/μL for all samples.

From all the potential pro-angiogenic targets we chose *Vegf* and *Hif-1a* as they are known to be the main drivers of vascular development in tumors. Additionally, *Dek* and *Stat3* were used as they influence *Vegf* expression in an *Hif-1a* independent manner^36,37^. To analyze the hypoxia-independent upregulation of *Hif-a* we used *Raptor*^38^. All primers were obtained from Integrated DNA Technologies (Coralville, Iowa, US) using sequences designed in-house that were BLASTed for specificity, and whose efficiency was determined using the standard curve method. Primer sequences, efficiencies, and corresponding annealing temperatures appear in Table S.1 in the Appendix.

The Luna Universal qPCR Master Mix (NEB, M3003) was used for qPCR employing 2 μL of cDNA sample and 18 μL mix of master mix (water, primers and qPCR mix) per well in 96-well plates (HSP9601, Bio-Rad), which were sealed using transparent film (MSB1001, Bio-Rad). All qPCR experiments were performed in a CFX96 Real Time PCR system (Bio-Rad) and were run in triplicate. Changes in the relative mRNA expression of all genes were quantified using Pfaffl’s method^39^ with murine glyceraldehyde-3-phosphate dehydrogenase (Gaphd) as the housekeeping gene^40^, and the expression of each gene in the CG group at week 1 as the normalizer.

### 3.6 Immunostaining

After the 6th (last) endoscopy for DRS and biopsy acquisition, mice were euthanized and colons were excised and cleaned using phosphate-buffered saline solution.Tumors were identified by comparing their anatomical location with the available imaging data in each case. Samples were stored in 10% neutral-buffered formalin for 24 hours, rinsed in saline, and later professionally embedded in paraffin and prepared by IHC World (Ellicott City, MD, US). Samples were sectioned in the sagittal plane to a thickness of 5 *μ*m prior to immunoperoxidase staining for Nestin-1 (Abcam, ab237036), a marker that is used to detect vessels in AOM-induced tumors^41,42^.

### 3.7 Image analysis

Tumor sections were imaged in an upright Nikon widefield microscope using a 10x objective (0.30 NA), and a gain of 1. Between 1-3 non-overlapping regions of interest (ROI) were obtained per sample, and in each ROI areas for segmentation were labeled as Nestin-positive (brown), background, or tissue (blue). The segmentation of each ROI was performed using the Trainable WEKA Segmentation from Fiji^43^. Between 1 to 4 regions of each label per ROI were manually selected and used to train the algorithm, followed by testing on the whole image. In each ROI, microvessel density (MVD) was quantified as the percentage of Nestin-positive area relative to total tissue area41. Additional details can be found in the Appendix.

### 3.8 Statistical Analyses

Statistical analyses were performed in R (4.1.1)^44^ using RStudio (2022.02.0 Build 333) and the {tidyverse}^45^ and {mgcv}^46,47^ packages. Specific information on the models used for each type of data is presented below.

#### 3.8.1 DRS and qPCR data

For the longitudinal data (DRS, qPCR) an assessment of the trends in the response over time showed that they did not follow a linear pattern (Figures 2, 3). Therefore we fitted generalized additive models (GAMs) with an interaction between time and treatment in each case. A GAM is a model that allows to fit non-linear trends in longitudinal data by using basis functions, thus overcoming the limitation of linear models (such as repeated measures ANOVA or a linear mixed model) which give biased estimates when used in data with non-linear trends^48^. Because GAMs allow the data to dictate the fit of the model, they are advantageous to perform comparisons between the different trends in the data. Pairwise comparisons between the different groups were obtained by calculating the difference in the fitted smooths in each case (the trend over time), and constructing a 95% simultaneous confidence interval (CI) around that difference^48,49^. In this way, the time intervals where a significant difference exists between the fitted trends will correspond to those intervals of time where the simultaneous CI does not cover zero^48,50^. Because the simultaneous CIs contain the whole function at a nominal level (95%), they are robust against false positives^47^. Additional details which include the syntax for GAMs for DRS and qPCR data, and figures with pairwise comparisons for all groups can be found in the Appendix.

**Figure 2:**
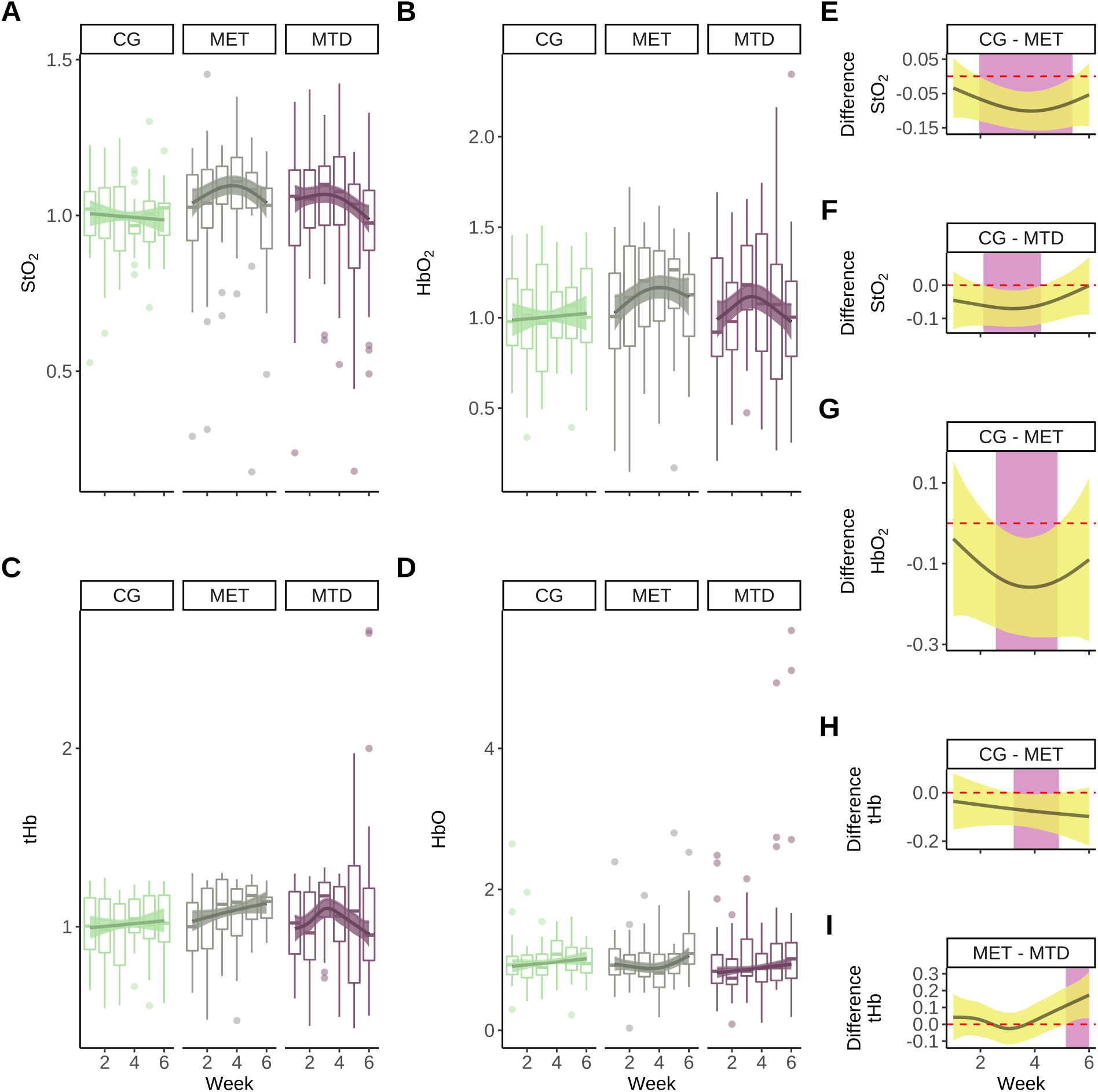
GAMs and pairwise comparisons for DRS-derived fold change values of perfusion. **A**: Longitudinal changes in StO_2_. The trend in the CG group remains flat while there is an increase that reaches a maximum at week 4 in the MET group and at week 3 in the MTD group. **B**: Longitudinal trends in HbO_2_, there is little change in the CG group whereas the MET and MTD groups show an increase that reaches it maximum at week 4 (MET) and week 3 (MTD). **C**: Changes in tHb, there is an increase in tHb in the MET and MTD groups over time. **D**: Changes in HbO, there is no change in the CG group over time, the MET and MTD groups show a temporal increase in HbO. In panels A-D lines represent the smooth trend fitted to the data and shaded regions represent a 95% simultaneous confidence interval. Boxplots show data distribution, points represent outliers. **E, F**: Pairwise comparisons of the smooth trends in StO_2_ shows a significant difference between the CG-MET and CG-MTD treated tumors between weeks 2 and 5 and 2 and 4, respectively. **G**: Pairwise comparisons of the smooth trends in HbO_2_ shows a significant difference between the CG and MET groups between weeks 3 and 5. **H, I**: There are significant differences in tHb between the CG and MET between weeks 3 and 4, and between the MET and MTD groups at week 5. In panels E-I the line represents the computed difference between the smooth trends in each group, shaded regions represent the 95% simultaneous confidence interval around the estimated difference, the purple area indicates time intervals of significance between the trends (where the confidence interval does not cover zero, which corresponds to the location of the red dotted line).

**Figure 3:**
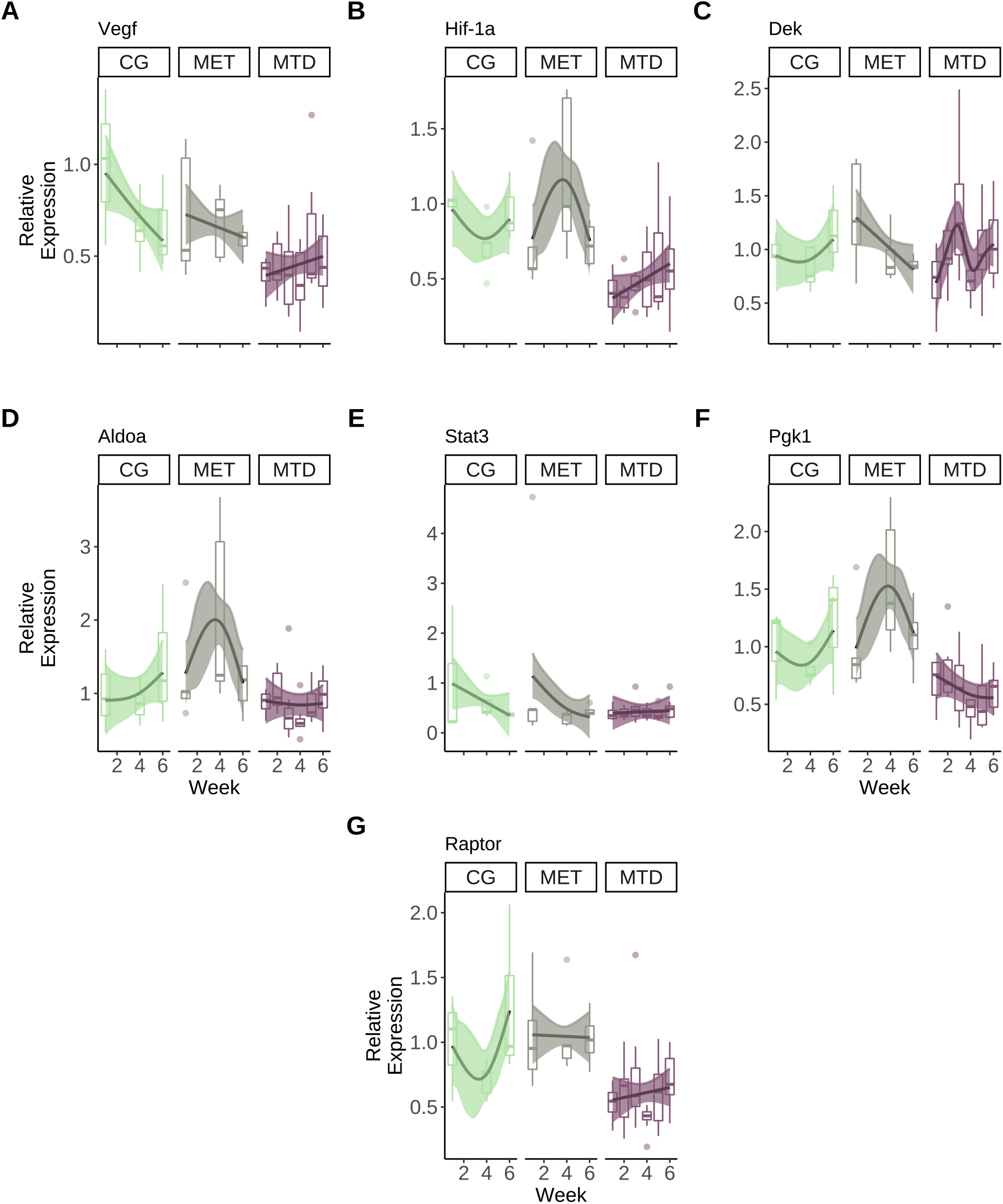
Longitudinal trends in the expression of different genes associated with perfusion in primary colorectal tumors. Boxplots present data distribution and points are outliers, lines represent the fitted smooth trend using the GAM. Shaded regions represent a 95% pointwise confidence interval around the smooth trend. **A**: Longitudinal trends in the expression of *Vegf*. **B**: Longitudinal changes in the expression of *Hif-1a*. **C**: Longitudinal changes in the expression of *Dek*. **D**: Longitudinal changes in the expression of *Aldoa.* **E**:Longitudinal changes in the expression of *Stat3.* **F**: Longitudinal changes in the expression of *Pgk1*. **G**: Longitudinal changes in the expression of *Raptor*.

#### 3.8.2 Imaging Data

For the imaging data, a Kruskal Wallis test was performed for the MVD values between groups to determine any statistically significant differences. Significant *p-values* were considered below 0.05.

## 4 Results

### 4.1 DRS-derived markers of perfusion

Using the endoscopically-acquired spectroscopy data we longitudinally quantified values of perfusion in all groups. For fold changes in StO_2_, the trend in the CG group showed no change across time, but an increase was observed in both the MET and MTD groups, with maximums of 8% at week 4, and of 5% at week 3, respectively (Figure 2A). For HbO_2_, the CG group showed no trend over time, and the MET and MTD groups showed an increase in the fold change value with a maximum mean rise of 15% and 10% at weeks 4 and 3, respectively (Figure 2B). Total hemoglobin (tHb) showed a similar trend in the three groups over time, where the mean increase reached a maximum of 13% and 11% in the MET and MTD groups at week 6 (Figure 2C). The trends for HbO showed no change in the CG group, and an increase over the six-week period in both the MET and MTD groups (Figure 2D) with mean maximum fold changes of 13% and 32%, respectively.

To determine if significant differences between the different groups existed, we performed pairwise comparisons to estimate the difference between the fitted smooths (via GAMs) for each DRS-derived perfusion value across the different groups. The rationale in this case is that when the difference between the fitted smooths is computed, a simultaneous confidence interval (s-CI) can be constructed around the difference, and the regions where the s-CI does not cover zero can be considered statistically significant periods of change. To improve visualization, we added a ribbon to highlight those periods where statistically significant differences existed. Significant differences existed in StO_2_ between the CG-MET and CG-MTD pairs. Specifically, between weeks 2 and 5 the MET group showed an increase in oxygenation (Figure 2E), whereas the increase was significant in the MTD group between weeks 2 and 4 (Figure 2F). For HbO_2_, the difference between the CG and MET groups was significant between weeks 3 and 4 (Figure 2G). For tHb, a significant differences existed between the CG-MET and the MET-MTD groups between weeks 3 and 5 and at week 5, respectively (Figure 2H, I).

### 4.2 Gene Expression

By obtaining repeated biopsies from tumors we were able to longitudinally examine the changes in expression in perfusion-related genes across the different treatment groups. Smooth trends (using GAMs) were obtained for each group in each case (Figure 3). Although the expression of *Vegf* showed a decrease over time in the CG and MET groups and an increase over time was observed in the MTD group, the MTD-treated tumors showed a 50% reduction in the expression of *Vegf* at week 1. Over the course of the study, the expression of *Vegf* in the MTD-treated group remained below the levels observed in the other groups (Figure 3A).

For *Hif-1a*, a decrease in expression was observed in the CG group between weeks 1-4 followed by an increase between weeks 4-6; a increase between weeks 1-4 followed by a decrease between weeks 4-6 was observed in the MET group; and a 50% reduction at week 1 was observed in the MTD group, followed by an increase over the six-week period (Figure 3B).

For *Dek*, an increase in the CG group between weeks 4-6 was observed, whereas the MET group showed a decrease over time. The MTD group showed a maximum increase at week 3, followed by a decrease between weeks 3-4 and an increase between weeks 4-6 (Figure 3C). However, the range of variation in all groups was similar.

Because HIF-1*α* is post-translationally regulated we also analyzed its downstream effects by analyzing the expression of *Aldoa* and *Pgk1* to indirectly assess its expression. For *Aldoa*, the expression over time showed an increase in the CG group, and a maximum was reached in the MET group at week 3; in the MTD group, there was no change in the trend over time and the range of the values was close to the expression of the gene at week 1 (Figure 3D).

In the case of *Pgk1*, the same trends from *Aldoa* were observed in the CG and MET groups, where the MTD group showed a reduction of ≈ 25% in expression at week 1 (3F).

In the case of *Stat3*, the trends in the three groups had a similar linear profile and showed minimal variation through the duration of the study (Figure 3E).

Finally, for *Raptor* a decrease in the expression between weeks 1-4 was observed in the CG group, followed by an increase between weeks 4-6. In the MET group, no change in the trend was observed, whereas a reduction of 50% was observed in the MTD group at week 1, followed by an increase through the duration of the study (Figure 3G).

Next, we performed pairwise comparisons to estimate the difference between the fitted trends for each group in each of the genes analyzed. For VEGF, there were significant differences between the CG-MTD and MET-MTD groups. The MTD group showed a significant reduction in *Vegf* expression of ≈ 60% at week 1, but the magnitude of the reduction diminished as the study progressed and became non-significant at week 4 (Figure 4A, B).

**Figure 4:**
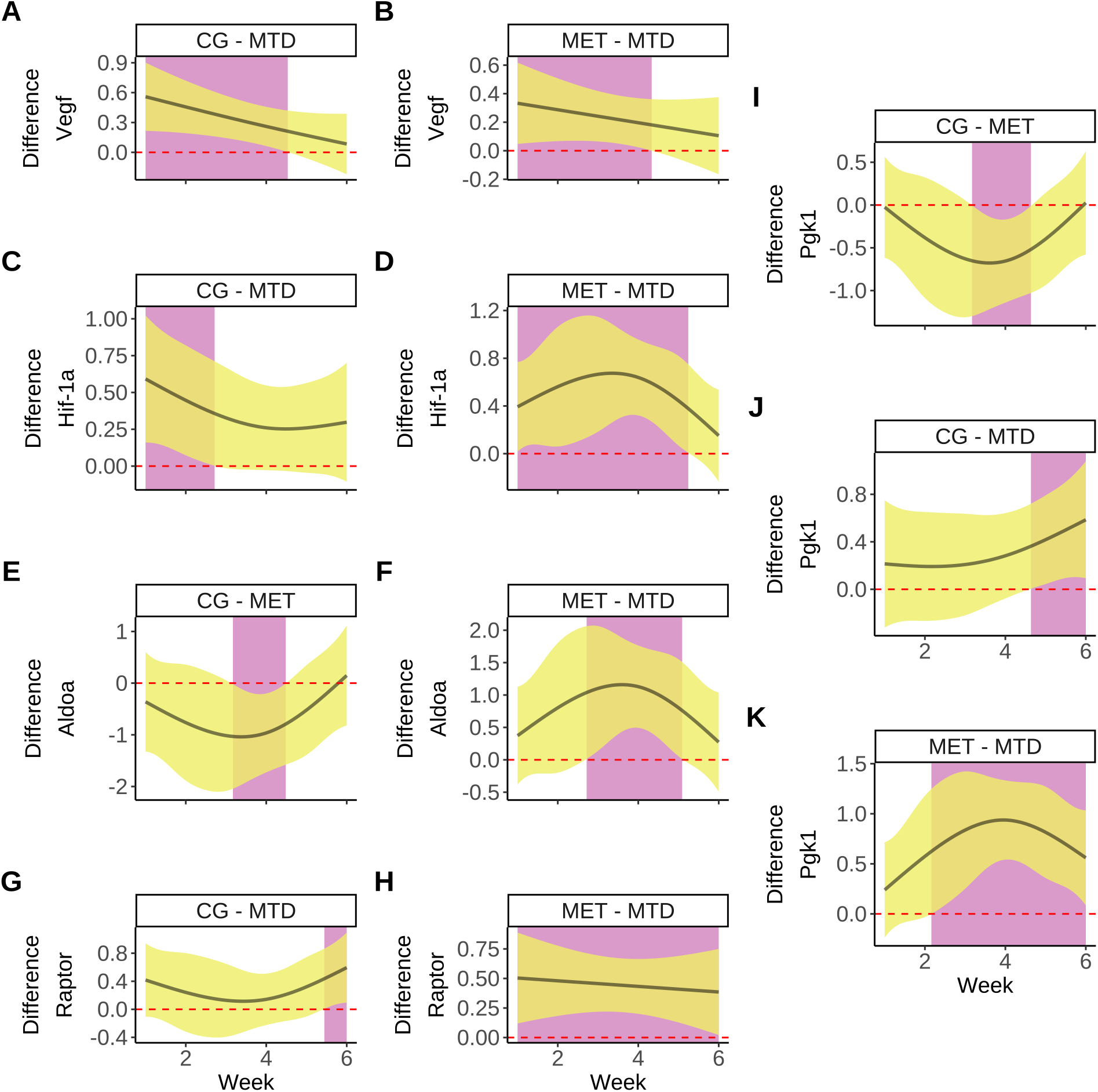
Pairwise comparisons between the fitted smooth trends for each group from Figure 3. Solid lines represent the estimated difference between the smooth trends, yellow shaded regions represent a 95% simultaneous confidence interval around the difference. Purple areas indicate time intervals where a significant difference exists (where the interval does not cover zero, which is marked by the red dotted line). In each panel, the first group of the pair is the one used as the reference to which the comparison of the other group is made. **A**, **B**: Significant differences exist in *Vegf* expression between the CG-MTD and MET-MTD groups between weeks 1-4.**C**, **D**: Significant differences exist in *Hif-1a* expression between the CG-MTD groups (weeks 1-2) and MET-MTD (weeks 1-5). **E**, **F**: Smooth trend comparisons for *Aldoa* expression show significant differences between the CG-MET (between weeks 3-4) and MET-MTD (weeks 3-5) groups. **G**, **H**: Significant differences in the longitudinal trends in expression for *Raptor* exist between the CG-MTD (week 5) and MET-MTD (weeks 1-6) pairs. **I**, **J**,**K**: The expression of *Pgk1* shows significant differences between the different groups. For the CG-MET pair the difference is from middle of week 3 through middle of week 4. In the CG-MTD pair, the difference is between weeks 5-6. In the MET-MTD pair, the difference is from week 2 until week 6.

For *Hif-1a*, significant differences exist between the CG-MTD groups from week 1 until the middle of week 2, and between the MET-MTD groups starting at week 1 until week 5 (Figure 4C, D). In both the CG and MET groups, the expression of *Hif-1a* was higher than in the MTD group by ≈ 50% at week 1, and the magnitude of the difference decreased as treatment progressed.

In the case of *Aldoa*, the overall difference in the trends followed the same pattern as those from *Hif-1a*, but the differences were significant between the CG and MET groups between weeks 3 and 4 (Figure 4E). In the case of the MET and MTD groups, significant differences existed between weeks 3 and 5 (Figure 4F). There were significant differences in the expression of *Raptor* between the CG and MTD (week 5), and MET-MTD (weeks 1-6) groups.

The pairwise comparison of the smooth trends for the relative expression of PGK1 showed the same trends from *Hif-1a* and *Aldoa*, but in this case significant differences existed between all groups. The expression of this gene was significantly lower in the MET group compared to the CG group between weeks 3-4 (Figure 4I). The CG group had higher expression than the MTD group between weeks 4 and 6 (Figure 4J). Finally, the expression in the MET group was significantly higher than in the MTD group between weeks 2 and 6 4K).

### 4.3 Effect of NAC in MVD in tumors

To determine the overall effect of both MET and MTD NAC in the development of vasculature in CRC, tumors were subjected to immunohistochemistry for Nestin in order to quantify the area of blood vessels relative to total tissue area, which represented the MVD of tumors. Specifically, MTD-treated tumors showed reduced vascular area (median 7.73% vs 25.1% and 23.63% in the CG and MET groups, respectively). There was a significant difference between the CG-MTD and MET-MTD groups in MVD (*p*=9.0 × 10^-7^ and *p*=3.9 × 10^-7^)(Figure 5D).

**Figure 5:**
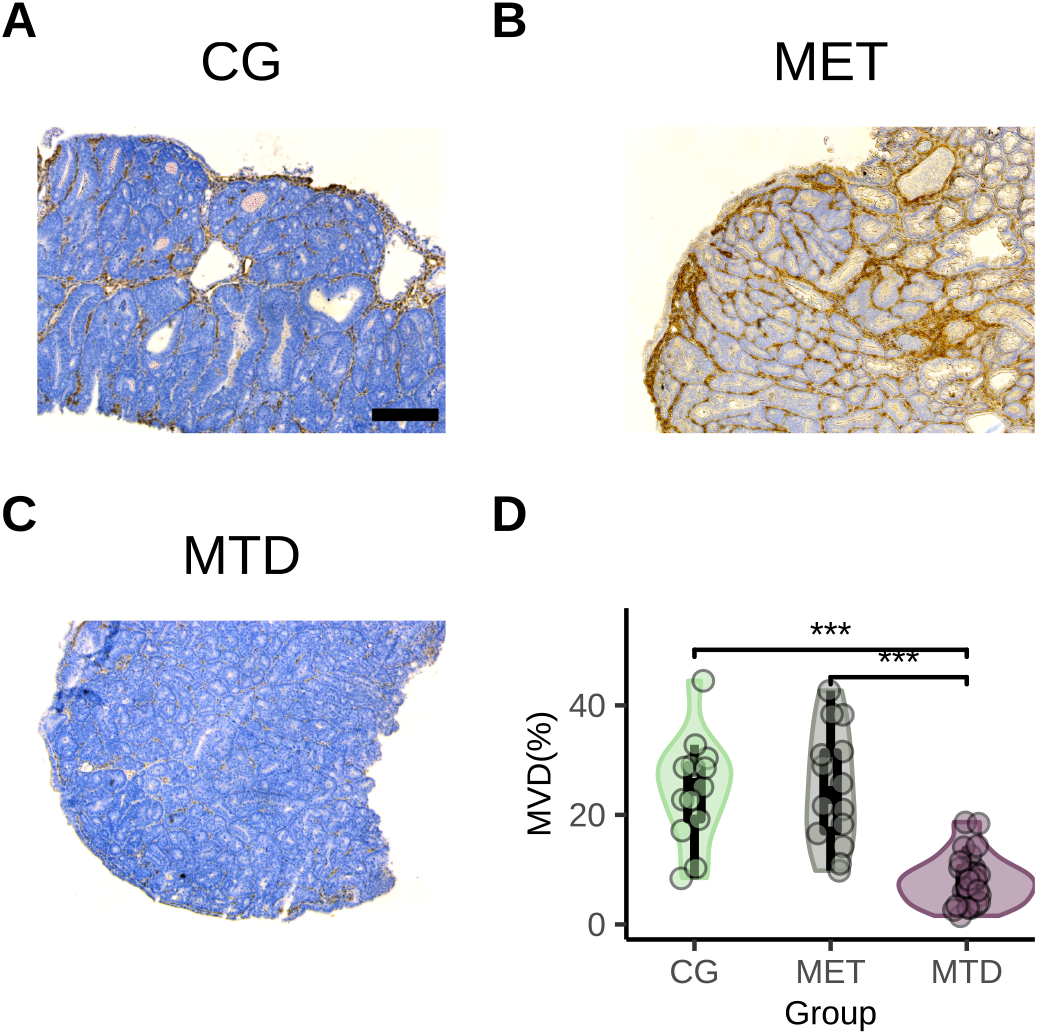
Immunohistochemical expression of Nestin in primary colorectal tumors treated with different NAC strategies. Nestin-positive regions (brown), Nestin-negative tissue (blue). **A**: Control (CG). **B** Metronomic (MET). **C**: Maximum-tolerated dose (MTD). **D**: Violin plot showing the density and spread of MVD in the different treatment groups. Significant differences exist for the total area of Nestin present between the CG-MTD (*p*=9.0 × 10^-7^) and MET-MTD-treated tumors (*p*=3.9 × 10^-7^) (Kruskal Wallis test). All images were obtained using a 10x objective, scale bar 200 *μ*m.

## 5 Discussion

Although MTD NAC in CRC aims to maximize the number of cancerous cells to be killed, this ultimate goal is limited by the rest periods it requires which lead to tumor re-growth and resistance^12,13^. In recent years, there has been an ongoing interest in the use of doses of cytotoxic drugs below the MTD threshold, which has led to the development of MET as an alternative treatment strategy for cancer. However, so far MET has not been explored as an alternative to MTD NAC in CRC. A critical component that needs to be addressed before MET NAC can be studied at the clinical level in CRC pertains its effect on the angiogenic program of tumors. Because tumor response is a microenvironment-dependent process, there is a need to examine if MET is able to cause the same anti-angiogenic effect in primary tumors that has been previously reported in subcutaneous models of this disease^51,52^.

In this regard, one way that tumor response to therapy (chemotherapy, radiotherapy or their combination) has been extensively studied in the past involves the use of DRS as a low-cost, non-destructive approach that quantifies changes in perfusion parameters (StO_2_, HbO_2_, etc.) which are known to change after therapy^25,27,53–56^. However, perfusion values that are quantifiable via DRS only represent ‘high level’ view of a preceding sequence of molecular shifts in the angiogenic program of tumors. Quantifying the changes in expression of genes involved in angiogenesis over the course of therapy may provide insight on the molecular mechanisms (‘low level’ changes) involved in the observed perfusion values obtained via DRS.

Specifically, because in the classical angiogenesis pathway the main driver of vascular development is *VEGF* (itself being upregulated by *HIF-1A* under hypoxic conditions^57^), we hypothesized that the use of endoscopic DRS and the longitudinal analysis of the levels of expression of *Vegf* and *Hif-1a* mRNA (and also the levels of mRNA from genes are known to upregulate *VEGF* and *HIF-1A* in an independent manner) would be able to provide a more comprehensive picture of the effect of MET and MTD NAC regimens in tumor angiogenesis in CRC. Our results suggest that while MET and MTD NAC elicit similar responses in perfusion, they cause distinct molecular responses in tumors under 5-FU chemotherapy. Considering the current interest in MET, our study points to the importance of the use of multimodal approaches that can provide insight on the different mechanisms that the angiogenic program of tumors undergo during therapy.

Our DRS results show that NAC (either MTD or MET) causes a similar effect in the trends of StO_2_ and HbO_2_ during the course of therapy, where an increase is observed during the first weeks of treatment followed by a gradual return to the baseline during the second half (Figure 2A, B). The estimated differences between the trends of StO_2_ indicate that in our model, MET causes an increase in oxygenation that can be detected via DRS and that becomes significant between weeks 2 and 5, whereas in the MTD the significant difference occurs between weeks 2 an 4 (Figure 2E and F, respectively). Although only the MET group showed a statistically significant increase in HbO_2_ when compared to the CG group between weeks 3 and 5 (Figure 2G), the same trend was observed for the CG-MTD pariwise comparison (Figure S.2,B in the Appendix).

The significant increase in StO_2_ we observed in the MET and MTD groups is consistent with the longitudinal elevation of this metric that has been reported in breast tumors, head and neck cancer xenografts and primary murine colorectal tumors after therapy^54–56^. In the past, the rationale for such increase has had multiple hypotheses: an increase in perfusion (when coupled with an increase in HbO_2_)^55,58,59^, the generation of free radicals by cytotoxic drugs that damage the nucleic acids of cancerous cells^60^, changes in metabolism^25^, and decreased oxygen consumption in surrounding tissues^59^. However, so far there have been no studies that provide longitudinal molecular information to those observed changes in perfusion. Here, we addressed the need of providing molecular context to the observed changes in perfusion by quantifying mRNA expression in genes associated with angiogenesis.

In this regard, we started by quantifying the expression of *Vegf* across all groups. Our analysis shows that both MET and MTD NAC are able to decrease the relative expression of *Vegf* when compared to the levels of untreated tumors, but such reduction is only significant for the MTD group through the first 4 weeks of treatment when compared to the MET and CG groups (Figure 4A, B) and can be measured 24-h after the first 5-FU injection. The average magnitude of the reduction is greater in the MTD group (≈ 60%) when compared with the MET group (≈ 29%). This would indicate that the increase in StO_2_ that is found via DRS in the MTD group (when compared to the CG group) through weeks 2 and 4 is the physiologically measurable outcome of the reduction in *Vegf* expression that starts at early at the beginning of the treatment phase, and that such increment in StO_2_ is likely to be driven by an alteration of the tumor vasculature due to therapy (an anti-angiogenic effect). Interestingly, because in the MET group the change in StO_2_ is not coupled with a significant change in *Vegf* expression, it appears that this type of therapy is able to affect tumor perfusion without a concomitant anti-angiogenic effect.

Additionally, because there was an endoscopically-visible reduction in tumor size between each therapy cycle in the MTD group (Figure 1C), the reduction in *Vegf* expression, and similar trends in the increase of StO_2_ and HbO_2_ would indicate a reduction in oxygen demand in surrounding tissue due to the number of tumor cells killed after each round of therapy driven by an anti-angiogenic effect in MTD NAC, consistent with results we have previously reported^27^. In other words, it appears that the magnitude of the decrease in *Vegf* expression that MTD NAC is able to achieve (and that is significantly lower than that achieved in the MET group) leads to significant blood vessel reduction/remodeling in the tumor when compared to the MET and CG groups (Figure 5C, D). Such result is consistent with the significant reduction of MVD in MTD-treated colorectal xenografts reported by Shi et. al.^61^.

In the case of MET NAC, our observed non-statistically significant reduction in *Vegf* expression using 5-FU in a MET regimen is in line with the results of a study by Fioravanti et. al. that addressed this same question31. Taken together, our results suggest that the observed changes in angiogenesis in the MET group are caused by an increase in perfusion, consistent with previous studies that have provided the same rationale to a simultaneous increases in both StO_2_ and HbO_2_ in tumors after therapy^55,58,59^. Furthermore, the changes in tHb through therapy show that in the MET group there is a significant increase between weeks 3 and 4 when compared to the CG group (Figure 2H), but that the MET group showed a significant increase in tHb at week 5 when compared to the MTD group (Figure 2I). Considering that by the end of therapy there is no significant difference in MVD between the MET and CG groups, but that the blood vessel area is significantly reduced in the MTD group (Figure 5D) this would support our previous analysis of the effects of each type of therapy in the angiogenic program of tumors. Specifically, there seems to be a distinct anti-angiogenic effect of MTD NAC in our model which leads to a remodeling of the tumor vasculature which results in reduced tHb. In contrast, the increase in tHb in the MET group seems not to come from an anti-angiogenic effect of therapy in the tumor vasculature and at the end of the treatment, there is no discernable difference between vessels in the CG and MET groups.

Additionally, the reduction in *Hif-1a* mRNA expression in the MTD group was significant when compared to the MET and CG groups (Figures 3B and 4C, D). Interestingly, in both NAC groups the mean reduction in *Hif-1a* expression at week 1 (≈ 25% and 60% for MET and MTD, respetively) closely matched the observed mean reduction in *Vegf* expression at the same timepoint. However, only in the MTD group the expression of both *Hif-1a* and *Vegf* followed the same overall trend (Figure 3A, B). Despite the fact that the connection between *Hif-1a* and *Vegf* to drive angiogenesis is well established^62^, there are no studies that analyze their expression in a longitudinal manner in tumors. In this case, at the very least, the similar levels of expression, longitudinal trends obtained for both genes, and the visually observable reduction in tumor size that MTD NAC causes suggest that in primary colorectal malignancies MTD NAC is able to effectively disrupt the VEGF-HIF-1a axis, thereby leading to a lasting anti-angiogenic effect that is not observable in MET NAC. However, it is well established that HIF-1A is post-transcriptionally regulated under hypoxia^62^, and therefore its effect is better quantified using downstream-regulated genes. To address this, we analyzed the levels of mRNA expression of two genes that encode enzymes involved in glycolysis and that are regulated by *Hif-1a*: *Aldoa (aldolase A*, one of the isoforms of the aldolase family)^63^ and *Pgk1* (*phosphoglycerate kinase 1*), which is one of the two ATP-generating enzymes in glycolysis^64^. Our results show that in the CG and MEG groups, the mRNA expression of *Hif-1a, Aldoa*, and *Pgk1* has a similar trend (Figure 3D, F). This result suggests that the majority of *Hif-1a* mRNA is not being post-transcriptionally degraded, thus being able to regulate *Aldoa* and *Pgk1* activity. Additionally, the pairwise comparisons for the smooth trends for both *Aldoa* and *Pgk1* show significant differences between the CG and MET groups between weeks 3 and 4 (Figure 4E, I), which is the time when the expression in the MET group is higher.

Interestingly, *Pgk1* and *Aldoa* do not show similar trends in expression with *Hif-1a* as seen in the MET and CG groups; in this case, there are significant differences between the MET and MTD groups during weeks 3 and 4 (*Aldoa*), and from week 2 until week 6 (*Pgk1*) (Figure 4F, K). This indicates the possibility of significant changes in metabolism caused by the dosing and frequency of therapy at the indicated time periods. It has been recently shown that in CRC cell lines, increased ALDOA expression enhances glycolysis by increasing glucose consumption and lactate production^65^, whereas in clinical samples PGK1 enhances proliferation^66^. Taken together, our results suggest that MET NAC enhances glycolysis and proliferation, whereas MTD NAC is able to suppress a glycolysis-driven metabolism in primary colorectal tumors. Additionally, the fact that increased *Hif-1a, Pgk1* and *Aldoa* expression are significantly different during extended time intervals when compared to the expression and smooth trends from the MTD group poses a question of the potential adverse effects of MET NAC in CRC. Because the efficacy of MET is dependent on the combined effect of drug dosing and frequency, our results indicate that a necessary step before the translation of MET NAC to the clinic involves drug dose and frequency optimization as it is possible that sub-optimal combinations are able to trigger resistance mechanisms (such as glycolysis upregulation) instead of limiting tumor growth.

Because angiogenesis is a complex process that does not rely exclusively on hypoxia to occur, we also analyzed three genes that regulate VEGF and HIF-1a by independent mechanisms. In the case of VEGF, we analyzed *Stat3* and *Dek*, which are known to regulate VEGF in a HIF-1a independent manner^36,37^. For *Hif-1a* we analyzed *Raptor*, as it encodes the RAPTOR protein which in turn regulates HIF-1a in an oxygen-independent manner^36,38,67^.

For *Raptor*, the trend in expression in the MET group remains relatively flat and similar in magnitude to that of the CG group, but the MTD group shows decreased expression at week 1 (≈ 50%) (Figure 3G). Moreover, significant differences exist in *Raptor* expression between the CG and MTD groups (between weeks 5 and 6), and the MET-MTD groups (beginning of week 1 until the end of the study) 4G, H). No significant differences were found between the pairwise smooth comparisons for *Stat3* and *Dek* across all treatment groups, indicating that both genes do not play a major role in regulating *Vegf* expression in primary colorectal tumors (Figure S.2 in the Appendix).

In the past, DRS has been used to quantify tumor response to therapy^27,58,59^, including the assessment of tumor response to MET in breast cancer^68^. Because DRS is minimally invasive, it provides information about early changes in perfusion after therapy, which can be used to evaluate the efficacy of different treatment strategies. However, because DRS is unable to provide metabolic or molecular information to the observed changes in perfusion, it is difficult to validate the hypotheses that can be inferred from such changes^25,55,60^. Traditionally, immunohistochemistry has been used to provide molecular context to DRS^59^, but this approach is limited as it is destructive and leads to a progressive reduction animal numbers in preclinical research which can in turn affect statistical power. In contrast, the analysis of changes in RNA expression in tumor biopsies can be used to obtain a sequential view of changes in tumor biology without the trade-off in the number of live specimens that immunohistochemistry presents. Our study shows that the analysis of mRNA expression can provide better context and support or refute hypotheses that can attempt to explain the molecular background of observed changes in perfusion via DRS. Specifically, it is interesting to note that in our model, similar trends in StO_2_ and HbO_2_ across different treatment strategies can have very different interpretations when analyzed in conjunction with mRNA expression.

On the other hand, the endosocopic application of DRS and the acquisition of colorectal tumor biopsies (where the placement of the instruments is done in an indirect manner) presents multiple challenges which can lead to artifacts in the collected signal due to pressure or probe misplacement^27^, or to obtaining biopsies or from non-matching sites. Despite this, we believe that due to their relative low cost and minimally invasive nautre, optical methods (such as DRS) can provide useful information of tumor biology when used longitudinally *in vivo* as there is potential in the use of the information generated by these technologies to determine their correlation with molecular markers of treatment response in CRC. In this way, DRS can be used to better to better understand the effects of therapy with regard to tumor resistance. Additionally, there is a need to obtain longitudinal information from tumors at the molecular level (using qPCR, RNA-seq, etc.) as such methods can provide powerful insight on the periods of increased cellular activity or changes in metabolism that can be exploited by novel therapeutic drugs and treatment strategies.

In summary, in this study we used a combination of diffuse reflectance spectroscopy and mRNA expression (via qPCR) to longitudinally assess changes in the angiogenic program of primary colorectal tumors under MTD and MET NAC regimens. In our model, MET and MTD NAC cause an increase in oxygen saturation in tumors that is quantifiable via DRS and becomes significant between the third and fourth week of treatment. When paired with the trends in mRNA expression, additional context can be provided to the observed changes in oxygenation. In the MTD group the increase in saturation is driven by reduced oxygen demand and is coupled with a significant reduction in *Vegf* expression when compared to tumors treated with MET NAC; such reduction leads to the remodeling of blood vessels, therefore causing decrease in MVD, which is quantifiable at the end of therapy. In MET-treated tumors the increase in oxygen saturation is caused by an increase in perfusion, and because *Aldoa* and *Pgk1* have increased expression, glycolysis appears to be upregulated in the MET group. Our study also demonstrates that when used in tandem, optical and molecular techniques can provide a comprehensive picture in the effects of novel treatment strategies. Our long term goal is to incorporate RNA-seq and optical techniques (DRS, multiphoton microscopy, fluorescence lifetime imaging) to holistically examine tumor biology and identify opportunities and challenges for novel therapeutic approaches in colorectal cancer. Future work will also incorporate the use of male mice to address potential sex-specific effects of chemotherapy.

## 6 Acknowledgments

This work was supported by the National Science Foundation CAREER Award (CBET 1751554, TJM), the Arkansas Biosciences Institute (TJM, AM), and the Fulbright Commission (Faculty Development Program, AM). The authors thank Regina Cabrera and Eleanor Schrems for their help with animal care, endoscopic procedures, and data acquisition.

## Supplementary Materials

An Appendix which contains qPCR methods, the GAMs used to fit longitudinal data and expanded figures for pairwise comparisons are available in PDF format. A repository that contains all the DRS and qPCR data, as well as the code used to create this manuscript can be found in https://github.com/aimundo/Primary_tumor_longitudinal_chemotherapy. The dataset of immunohistochemistry images is stored in Biostudies under accession number S-BIAD323.

