## Appendix for "Longitudinal examination of perfusion and angiogenesis markers in primary colorectal tumors shows distinct signatures for metronomic and maximum-tolerated dose strategies"

SUPPLEMENTARY MATERIALS: APPENDIX

### S MATERIALS AND METHODS

#### S.1 Primers

Primer sequences were designed in-house, BLASTed for specificity and obtained from Integrated DNA Technologies (Coralville, Iowa, US). Due to the relatively low amount of cDNA obtained from each tumor sample we used CT-26 cell derived cDNA for efficiency and annealing temperature determination via the standard curve method using 6 serial dilutions (5x each). All experiments were performed on a CFX96 Real Time PCR system (Bio-Rad).

Primer information regarding sequences, annealing temperatures, and efficiencies can be found in Table S.1.

Table S.1: Primer information

| **GENE** | **SEQUENCE (F/R)** | **PRODUCT LENGTH (bp)** | **ANNEALING TEMP. °C** | **AMPLIFICATION EFFICIENCY (%)** |
|---|---|---|---|---|
| GAPDH | CCTCGTCCCGTAGACAAAATG/ TGAAGGGGTCGTTGATGGC | 124 | 60.0 | 96.25 |
| VEGF | TAACGATGAAGCCCTGGAGT/ TATGTGCTGGCTTTGGTGAG | 91 | 60.0 | 97.85 |
| HIF-1a | CCTGATGCTCTCACTCTGCT/ TTCAAGTTGTTGATCTTCAGTTTC | 99 | 60.0 | 101.72 |
| DEK | GCAGACAGCAGTACCACCAA/ TCATCTGTAGGTGGCTTTTTCA | 116 | 60.0 | 98.61 |
| PGK1 | AGAGTCCAGAGCGACCCTTC/ TTATTGATCAGCTGGATCTTGTC | 73 | 55.7 | 91.99 |
| RPTOR | TGGTGAAGACCACACCCTGT/ GTAGGTTTGCACCGATGGTT | 96 | 60.0 | 99.42 |
| ALDOA | ACGGTCACAGCACTTCGTC/ TTGATGGATGCCTCTTCCTC | 92 | 64.5 | 97.43 |
| STAT3 | GAGGGGTCACTTTCACTTGG/ TCAGCTGCTGCTTGGTGTAT | 87 | 58.0 | 93.90 |

#### S.2 Statistical Analyses

##### S.2.1 Generalized Additive Models (GAMs)

For both the DRS and qPCR data a GAM to analyze the trend of the variable of interest (either DRS-derived pefusion parameters or relative gene expression) was fitted, using the package `{mgcv}`. The overall construction of the model was:

```
model<-gam(Variable ~ Group +s(Day, by = Group, k=6, method = 'REML', data = data)
```

Where `model` specifies the object that stores the fitted model, `Variable` represents the variable of interest (qPCR fold changes or DRS-derived optical parameters), `Group` is a parametric term that allows to provide an intercept to each of the three treatment groups, and the content of the smoother (starting with `s()` specifies the use of 6 basis functions (`k=6`) where the smoothness parameter is estimated via restricted maximum likelihood (`method=REML`). Finally, `data` refers to the dataframe containing the data of interest.

**S.2.1.1 DRS** The next chunk shows all the models fitted for the DRS data.

```
StO2_gam <- gam(StO2_fold ~ GROUP+s(WEEK, by = GROUP, k = 6),
               family=scat(link='identity', min.df = 3),
               method='REML',
               data   = data1)

HbO2_gam <- gam(HbO2_fold ~ GROUP+s(WEEK, by = GROUP, k = 6),
               family=scat(link='identity', min.df = 3),
```

```
                  method='REML',
                  data   = data1)

tHb_gam <- gam(tHb_fold ~ GROUP+s(WEEK, by = GROUP, k = 6),
                 family=scat(link='identity', min.df = 3),
                 method='REML',
                 data   = data1)

HbO_gam <- gam(HbO_fold ~ GROUP+s(WEEK, by = GROUP, k = 6),
                 family=scat(link='identity', min.df = 3),
                 method='REML',
                 data   = data1)
```

The code to obtain Figure 3 on the main manuscript (where the individual values are shown along with the models) can be found in the script DRS_GAMs.R

Figure S.1 is an expanded version of the pairwise comparisons from Figure 2, showing all the pairwise comparisons between the different DRS-derived measures of perfusion.

**S.2.1.2 qPCR** Here, we present the GAMs used to fit the trends in mRNA expression, and which can be found in the script qPCR_GAMs.R, which also contains all the previous steps for data cleaning, normalization, and relative expression calculation.

```
k=6
VEGF_gam<-gam(Norm_ratio~GROUP+s(WEEK,by=GROUP,k=k),
              method='REML',
              data=VEGF)

HIF1_gam<-gam(Norm_ratio~GROUP+s(WEEK,by=GROUP,k=k),
              method='REML',
              data=HIF_1)

DEK_gam<-gam(Norm_ratio~GROUP+s(WEEK,by=GROUP,k=k),
             method='REML',
             data=DEK)

ALDOA_gam<-gam(Norm_ratio~GROUP+s(WEEK,by=GROUP,k=k),
               method='REML',
               data=ALDOA)

PGK1_gam<-gam(Norm_ratio~GROUP+s(WEEK,by=GROUP,k=k),
              method='REML',
              data=PGK1)

RPTOR_gam<-gam(Norm_ratio~GROUP+s(WEEK,by=GROUP,k=k),
               method='REML',
               data=RPTOR)

STAT3_gam<-gam(Norm_ratio~GROUP+s(WEEK,by=GROUP,k=k),
               method='REML',
               data=STAT3)
```

Next, Figure S.2 is an expanded version of Figure 4 from the main manuscript. It presents all the pairwise comparisons between the fitted smooths for all genes. Only the pairwise comparisons with significant differences (interval not covering zero), are included in Figure 4.

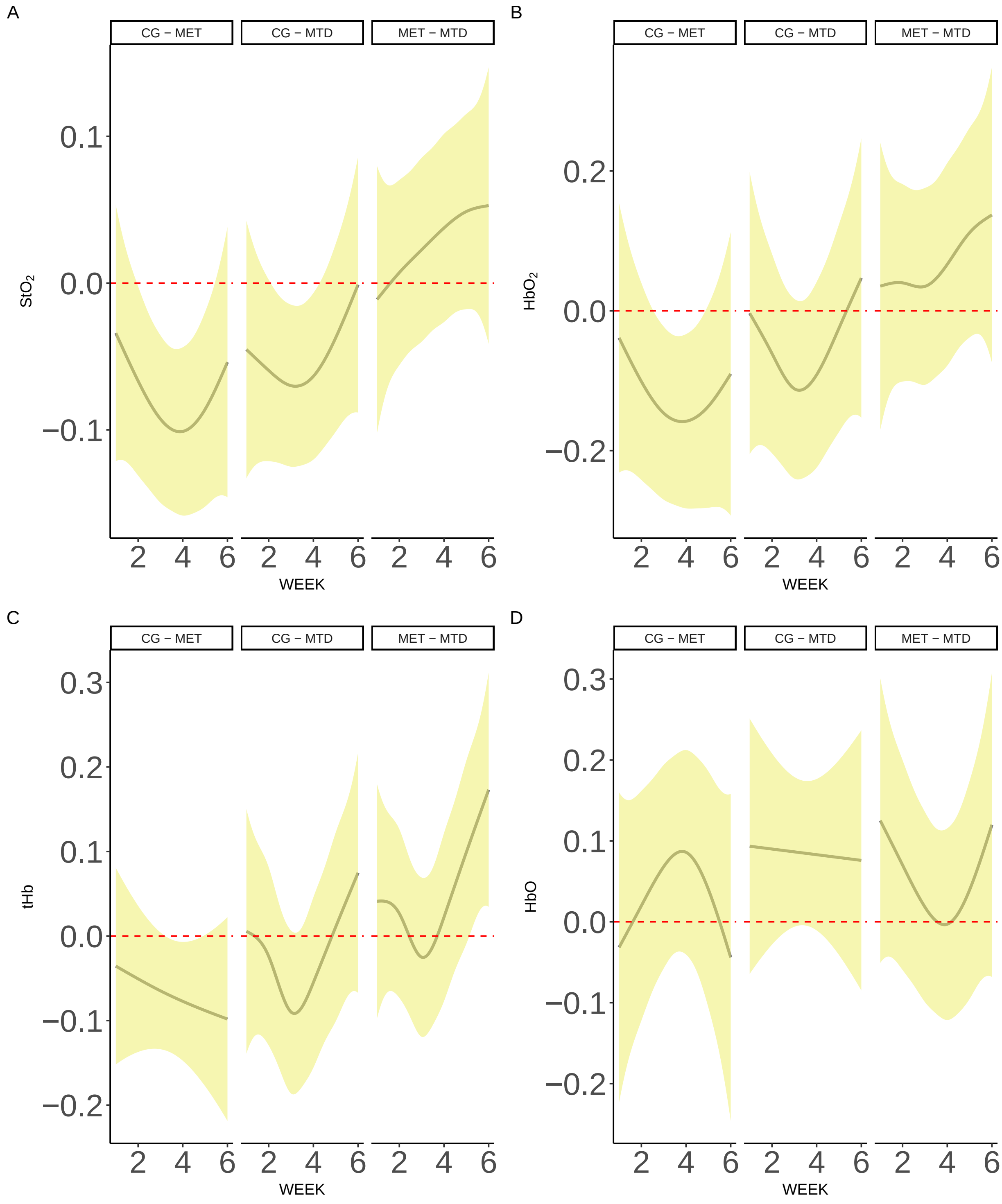


Figure S.1: Smooth difference pairwise comparisons for all DRS-derived data.

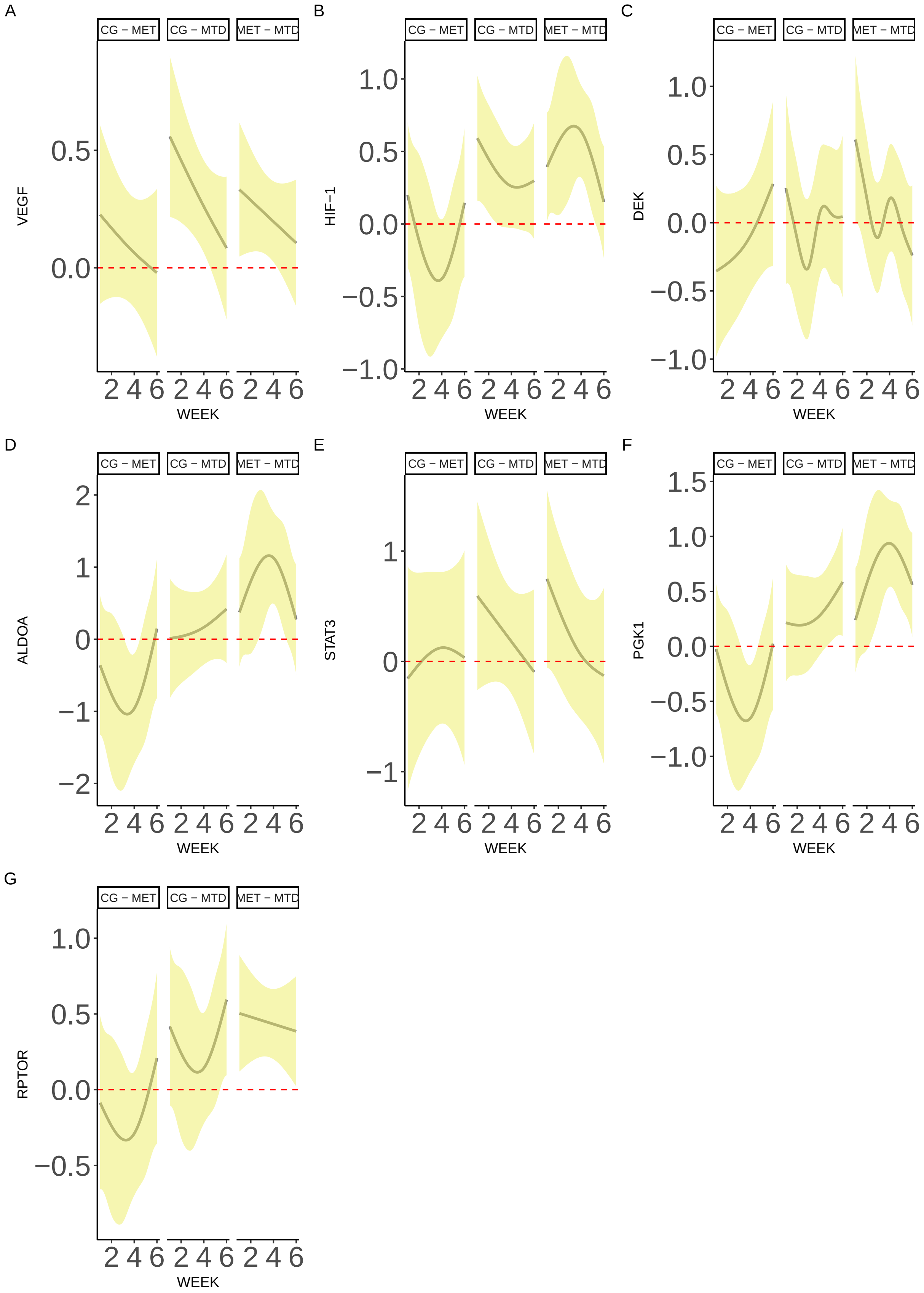


Figure S.2: Smooth difference pairwise comparisons for mRNA expression across all genes.

#### S.3 Image Analysis and MVD Quantification

Tumor sections were imaged in an upright Nikon widefield microscope using a 10x objective (0.30 NA) and the same binning, exposure time and gain settings. Between 1-3 non-overlapping regions of interest (ROI) were obtained per sample, and in each ROI areas for segmentation were labeled as Nestin-positive, background, or tissue. The segmentation of each ROI was performed using the Trainable WEKA Segmentation from Fiji. Between 1 to 4 regions of each label per ROI were manually selected and used to train the algorithm, followed by testing on the whole image. Figure (ref:fig-MVD-Appendix)A shows the image analysis workflow. Figure (ref:fig-MVD-Appendix)B shows a representative tumor image, where the Nestin-1 positive staining is visible (brown areas). Once the ROIs were labeled accordingly, the WEKA algorithm was used to obtain segmented regions (Figure (ref:fig-MVD-Appendix)C). Pixels identified as background (red regions in Figure (ref:fig-MVD-Appendix)C) were not considered for MVD quantification.

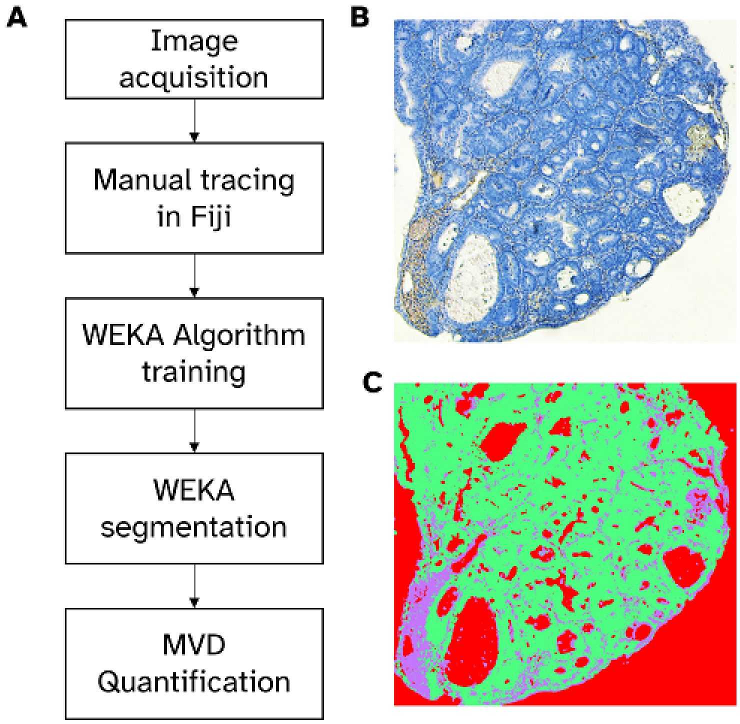


Figure S.3: MVD quantification workflow. **A**: Workflow for MVD quantification. **B**: Representative tumor image showing Nestin-1 positive staining (brown), negative Nestin-1 tissue staining (blue) and image background (white). **C**: Mask generated from **B** using the WEKA Algorithm. Nestin-1 positive areas (blue), tissue (green), background (red).
